# Variance Reducing and Noise Correction in Protein Quantification by Measuring Fluctuations in Fluorescence due to Photobleaching

**DOI:** 10.1101/2021.06.10.447938

**Authors:** Itay Gelber

## Abstract

Quantifying protein number using the ratio between the variance and the mean of the protein distribution is a straightforward calibration method in the experimental conditions for microscopy imaging. Recently the model has been expanded to decaying processes with binomial distribution. In this paper, we examine the model proposed, and show how the algorithm can be adapted to the case of variance in the initial number of proteins between cells. We propose improving the algorithm so that the information processing of each frame is done independently from other frames. By doing so, the variance in the process of determining the protein number can be reduced. In addition, we examine the handling of unwanted noises in the measurement, offer a solution for shot noise and background noise, and examine the expected error caused in calculating the decay constant. We also analyze the expected difficulties in conducting a practical experiment, which includes non-exponential decay, and variance in the decay constants of the cells. These methods can be applied to any superposition of *n*_0_ discrete decaying processes. However, the evaluation of expected errors in quantification is essential for early planning of the experimental conditions, and for the evaluation of the error.

## INTRODUCTION

Knowing the absolute number of proteins involved in a biological process is important for understanding process variability, nonlinearity, and comparing results from different laboratories. However, the contribution of a single protein to a measured intensity usually cannot be directly identified. In several cases, attempts have been made to address the problem by quantifying protein number using the ratio between the variance and the mean of the protein distribution, which provides a straightforward calibration for time-lapse imaging.

### Quantification via Calculation of the Ratio between Variance and Mean

In a system where many independent events occur, when the contribution to a single event signal cannot be directly measured, due to low sensitivity, the contribution of a single event can be calculated using the ratio of variance to mean measured value. That is, consider a system where *I_j_* = *αn_j_, I_j_* is the signal measured in measurement *j*. *n_j_* is the number of random events that occur in measurement *j*, and *α* - the contribution of each event to the measured signal.

In the specific case of a normally distributed number of random events (for which *σ*^2^ = *n*), the signal is 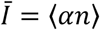 and the variance of the signal is *Var(I)* = *α*^2^ · *Var(n)* = *α*^2^*n*, the ratio between the variance and the mean gives *α*, the calibration parameter:

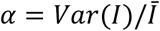

with which the absolute number of events can be calculated. This method is used, for example, to calculate the electron charge, by measuring the mean and variance of the number of electrons emitted from a cathode, independently of each other ([1]). Similarly, it used to measure the fractional charge in Fractional Quantum Hall Effect ([2], [3]). In the field of microscopy, it can be used in the calculation of the signal emitted from a single fluorophore, through the distribution of fluorescent proteins between two daughter cells during the division process([4], [5]). In [6] – in helps in the calculation of the ratio between the variance in fluorescence of the nucleoids, resulting from labeling the proteins attached to the nucleus, and their mean fluorescence, used to estimates the lower bounds for the number of proteins. The reason the lower bounds can only be estimated is the other factors in the formation of the variables. One such factor is mRNA fluctuations in space, or time, that affect the variance of the nucleoids. The advantage of this technique is that it does not require sensitivity to a single molecule, as in the case of stepwise photobleaching ([7], [8]). It is important to calibrate the fluorescence in vivo, under the original conditions of the laboratory experiment, since the fluorescence is affected by solvent polarity ([9]), pH ([10], [11]), or temperature ([12], [13]).

Rutenberg ([14]) expanded the model to decaying processes with binomial distribution. In this case, *σ*^2^ = *np*(1 - *p*), where *p* is the discrete probability distribution of the number of successes in a sequence of *n* independent experiments. Rutenberg proposed using the photobleach process of a fluorescent protein – a random process in which a fluorescent protein, under illumination, irreversibly loses its ability to emit light – to calibrate the signal from a single fluorophore, as measured by a microscope camera. He shows this method has a smaller variance than the methods described above, that it is faster, and is suitable for a wider range of cells.

with this method, cells that contain fluorescent proteins are placed under strong illumination so they undergo photobleaching. Throughout the process images of the fluorescence are taken continuously, using the microscope camera. Rotenberg notes that their analysis of photobleaching fluctuations can apply to any superposition of *n*_0_ discrete decaying processes.

Since the publication calibrations were made using this method to myosin Va molecular motor ensembles ([15], [16]) Venus-TnpA molecules ([17]), GFP molecules for Ter4 and Ori2 in Escherichia coli ([18]).

In this paper, we examine the model proposed by Rothenberg, and show how the algorithm can be adapted to the case where there is a variance in the initial number of proteins between cells. We also propose to improve the algorithm so that the processing of information obtained from each frame will be done independently from that of information from the other frames, and show that by doing so, the variance in determining the protein numbers can be reduced. In addition, we examine the handling of unwanted noises in the measurement, offer a solution for shot noises, and background noises, and examine the error expected to be caused by calculating the decay constant.

Rutenberg’s proposed theoretical model assumes Fluorescence decay is exponential. However, an article published later, by peter S. Swain ([19]), claims the decay curve of fluorescence due to photobleaching is apparently multiexponential due to variability in the environment the proteins are in within the cell, and in their orientation for light. He therefore proposes a model in which the expected decay is calculated using Gaussian Process ([20]), where it is not necessary to assume a specific decay model for quantification. He tested six proteins in yeast and showed their estimates of numbers agree, within an order of magnitude, with published biochemical measurements. The use of the Gaussian process technique has an advantage in that it is suitable for any model of decay. Its disadvantage, on the other hand, is that it allows for over-fitting, which leads to a reduction in the calculated noise, and an overestimation of the number of proteins.

### Quantification from Photobleach Fluctuations

In the method developed by Rotenberg, the random process is a photobleach of a fluorescent protein: a random process in which a fluorescent protein under illumination irreversibly loses its ability to emit light. If we define the average fraction of surviving fluorophores at time *t* using *p*, and assume all the fluorescent proteins are in equal environmental states, the following ratio will be obtained for uniform and constant illumination:

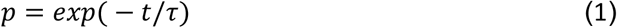

For many fluorophores, the average number that have not undergone photobleaching after time *t* is *n* = *n*_0_*p*, where *n*_0_ is the initial protein number. The variance in the number of proteins that survive depends on *p* and is: (binomial distribution)

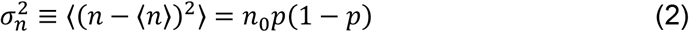

The signal measured by the camera can be expressed by:

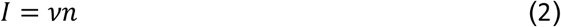

Where *I* is the light intensity measured from *n* fluorophores, and *ν* is the calibration factor, that determines the signal from a single fluorophore. Its value depends on illuminance, measurement time, system optics, fluorophore properties, and camera properties.

With the help of (2) and (3) it is possible to obtain an expression for the variance in the signal:

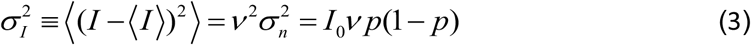

where *I*_0_ is the initial illumination. Now *ν* can be written:

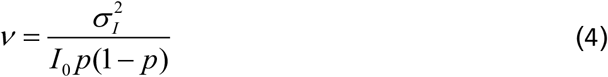

For calculating *ν*, cells containing fluorescent proteins are exposed to permanent illumination, under a microscope which causes photobleach. In the process it takes pictures continuously, so the illumination of each cell is measured over time. Monitoring the decaying fluorescence intensity of many cells makes it possible to calculate the decay constant τ (according to (1)). Since the time intervals between images are constant (T), the average fraction of surviving fluorophores after a single frame is defined as *p_f_* = *exp(− T/τ)*. Hence, the value *p* of frame *j* (*j* = 0,1,2 …) is 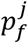. Rothenberg proposed taking the average of the fluorescence at any given time 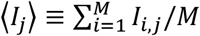, where *M* is the number of cells, *j* the number of the frame. He used it to calculate the variance of the fluorescence of the cells in frame *j*. Now, with the help of (5), we can get *ν*. Calculating and averaging *ν* for many cells leads to an accurate result. Rothenberg showed that the variance in the calculated result of *ν* from a single frame in a single cell, according to (5), is:

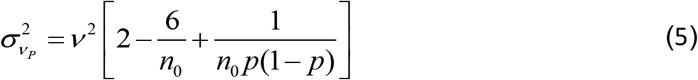

He proposed to make an average of the values obtained from each individual cell, by performing integration from *p* = 0 to *p* = 1 to both sides of Eq. (5) (if the measurement is made for a different *p* domain the limits of integration can be corrected), so that the following expression is obtained:

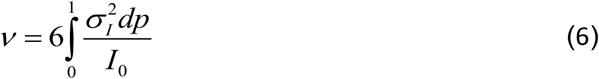

In this method the variance is given by:

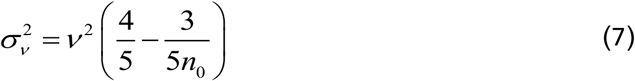

### Adapting the Method to the Case of Variance in the Initial Number of Fluorophores

An experimental ensemble of cells will have a distribution of initial fluorescence intensities [21]. Rotenberg argued that because the variance in the fluorescence, calculated for each cell, is divided by the initial fluorescence intensities of that cell(*I*_0_), the initial variance in the number of fluorophores does not cause an error. This claim relates to the calibration of the variance, with respect to the initial fluorescence of each cell. However, there is still an error in the calculation of the variance itself. For cell *i*, having an initial fluorescence different from the mean in the first frame, *I*_(0,*i*)_ ≠ ⟨*I*_0_⟩, the average fluorescence of the cells in frame *j* is not the value around which the probability *I_(j,i)_* is distributed. If *I*_(0,*i*)_ > ⟨*I*_0_⟩ then the measured intensity of *I_i_*, throughout all frames, is expected to be greater than ⟨*I_j_*⟩, and vice versa if it is small. Therefore, calculating the variance by 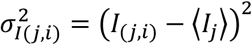 leads to an increase in measured variance. This error can be calculated by setting α to be the difference from the mean, so that ⟨*I*_0_⟩ = *I*_(0,*i*)_(1 + *α*). The expression for the error in calculating the variance, is 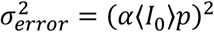 (Sup. 1). Whether the initial protein number is smaller or larger than the average, the result is an increase in the variance measured. Hence, the error will not be eliminated by averaging the results obtained from many cells. It can be shown that the relative error in the final calculation of *v* according to Eq. (7) (Sup. 1) is:

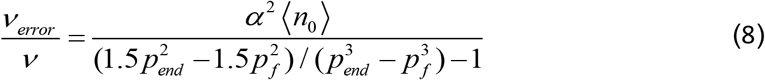

Where *p_end_* is the integral limit.

To avoid this error, which may be critical (Fig. 1), the variance in *p* values should be calculated when using Eq. (5) instead of the variance in fluorescence. If *p_(j,i)_* = *I_(j,i)_*/*I*_(0,*i*)_ then the variance in *p* values is given by 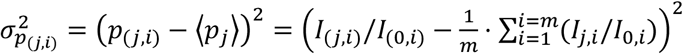.

**Figure 1:**
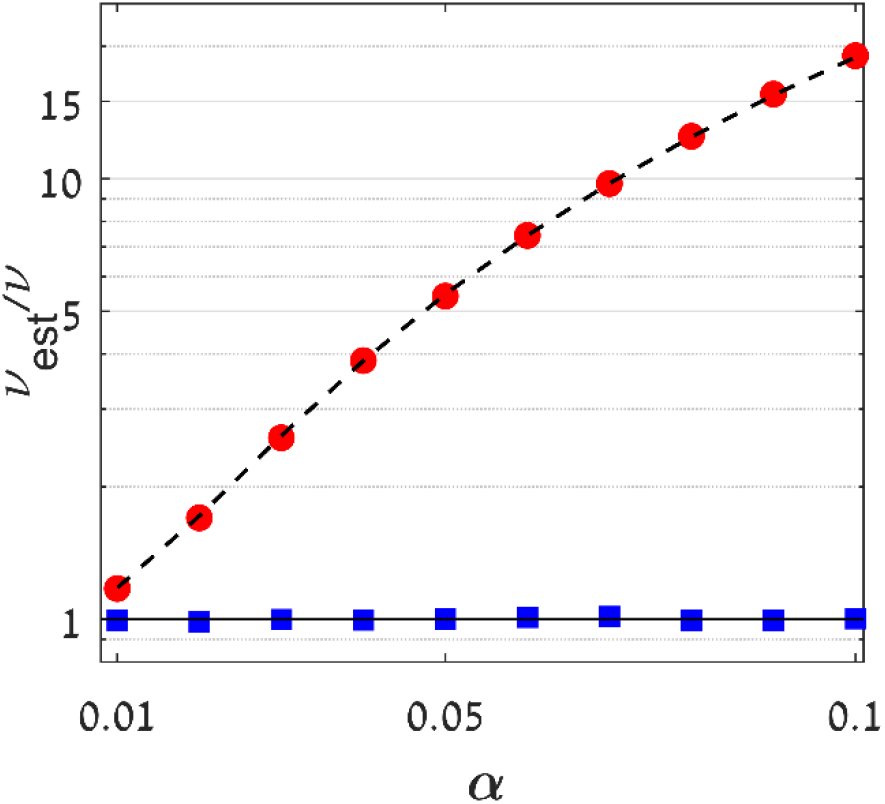
Simulation results showing *ν_est_*/*ν* vs. *α*. Red circles: Results according to Eq. (7). Black dashed line: Expected result according to (9). Blue squares: Results according to Eq. (10). For the simulation we used *p_end_* = 0.1, *n*_0_ = 1000, *p_f_* = 0.9, ν = 753 (According to the value used by Rotenberg). the. Calculations were done on 10^4^ cells.

This variance is not affected by the initial fluorescence values because each value of *I_(j,i)_* is calibrated relative to *I*_(0,*i*)_. since 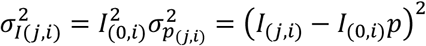 Eq. (5) can be use as follows:

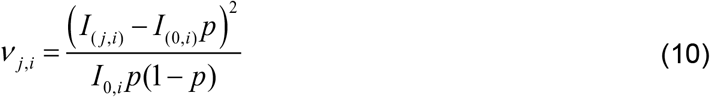

That is, Eq. (10) calculates the variance in frame j in relation to the value expected from the first frame (*I*_(0,*i*)_*p*). This as opposed to calculating the variance with respect to the average fluorescence of the cells, as in the method proposed by Rotenberg, without considering that the fluorescence from each cell is not expected to be distributed around the same value. Another advantage of calculating with respect to the initial value is that since p can be calculated from any pair of frames (*p_f_* = *I_j-1_/I_j_*), it is possible to eventually calculate it from an average of *(N – 1)M* values (where *N* is the number of frames, and *M* the number of cells), as opposed to calculating ⟨*I_(j)_*⟩ from *M* values.

For testing equations (9) and (10), a simulation was performed in which values of *ν_est_/ν* are calculated against *α* in both methods. For each *α* value half of the cells are with initial fluorescence *I*_(0,*i*)_ = ⟨*I*_0_⟩(1 + *α*), and half have *I*_(0,*i*)_ = ⟨*I*_0_⟩(1 − *α*). The effect of photobleaching is determined randomly for each fluorescent protein, for every frame. The value *p_f_* = 0.9 represents the probability of the protein surviving per exposure time of single frame. The simulation was performed on 10^4^ cells and the mean *ν* was obtained by measuring the variance in relation to the fluorescence of the cells in the same frame (red circles. The black dashed line represents a theoretical calculation according to Eq. (9)). The variance is measured in relation to the expected value from the first frame of the same cell (Eq. (10)) (blue squares). The results show that differences in the initial illumination values have a dramatic effect on calculating *ν*, using Eq. (7), and this effect is predicted exactly by Eq. (9). Be it as it may, Eq. (10) produces an accurate, unaffected result from the variance in initial values. In case there is no difference in the *n_0_* values between the cell’s, Eq. (5) and (10) are identical, so that throughout the rest of the article we refer only to Eq. 10.

### Calculating the Variance of Events Since the First Frame vs Previous Frame

Until now, *p* has always been in relation to the first frame (*I*_0_) (First Frame Comparison -FFC). Since the variance in each frame is due to all the random events in the previous frames, though, it is correlated with the variance measured in the previous frames. The final *ν* is calculated using the averaging of 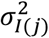, calculated from each frame. The variance of a sum of two random variables is given by *Var(X + Y)* = *Var(X)* + *Var(Y)* + 2 *Cov(X, Y)*. So, since 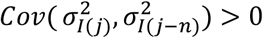, the final variance increases. To avoid this, the expected *I_j_* can be calculated separately from *I*_0_, but in relation to *I*_*j*-1_ (Previous Frame Comparison - PFC) i.e.:

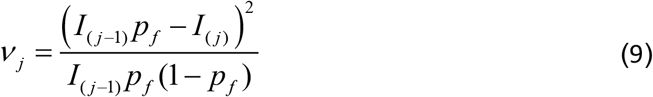

In this way, there is no correlation between the measurements from different frames, and therefore the final variance is smaller. In fact, according to Eq. (6), each measurement of a frame with respect to the previous frame has a variance of 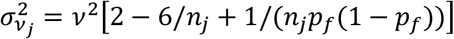. For *n_j_* > 100, the expression can be written as 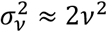. The final variance is obtained from the average of the results of all frames:

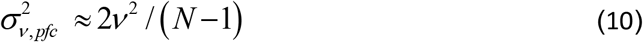

where *N* is the number of frames (so *N* - 1 variance values can be calculated). If we approach the variance obtained by (8) in the Rothenberg method, for *n*_0_ > 100, so 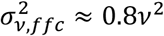, we can see that for *N* > 4, a lower variance is obtained by the PFC method.

The number of frames in the measurement is calculated using the definition *p_end_* – the ratio between the final signal, for which the measurement is stopped, for reasons of signal-to-noise ratio, and the initial signal. So, for example, if the measurement stops when the signal reaches 5% of the initial signal, i.e., 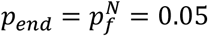, then *N* = *log_pf_*(0.05). Figure (2) shows 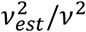 and 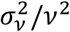 as calculated using a numerical simulation, according to the two methods. *ν* was calculated from simulation data for a single cell. For the purpose of averaging, 10^6^ measurements were made. 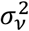 was also measured. The results are presented as dependent on *p_end_*. For the simulation, the parameters *n*_0_ = 1000,*ν* = 753,*p_f_* = 0.9 were used. The two methods calculate *ν* with great accuracy, without bias, but the variance obtained for FFC is significantly higher than for PFC. The results correspond exactly to the variance predicted by Eq. (8) and (12).

**Figure 2:**
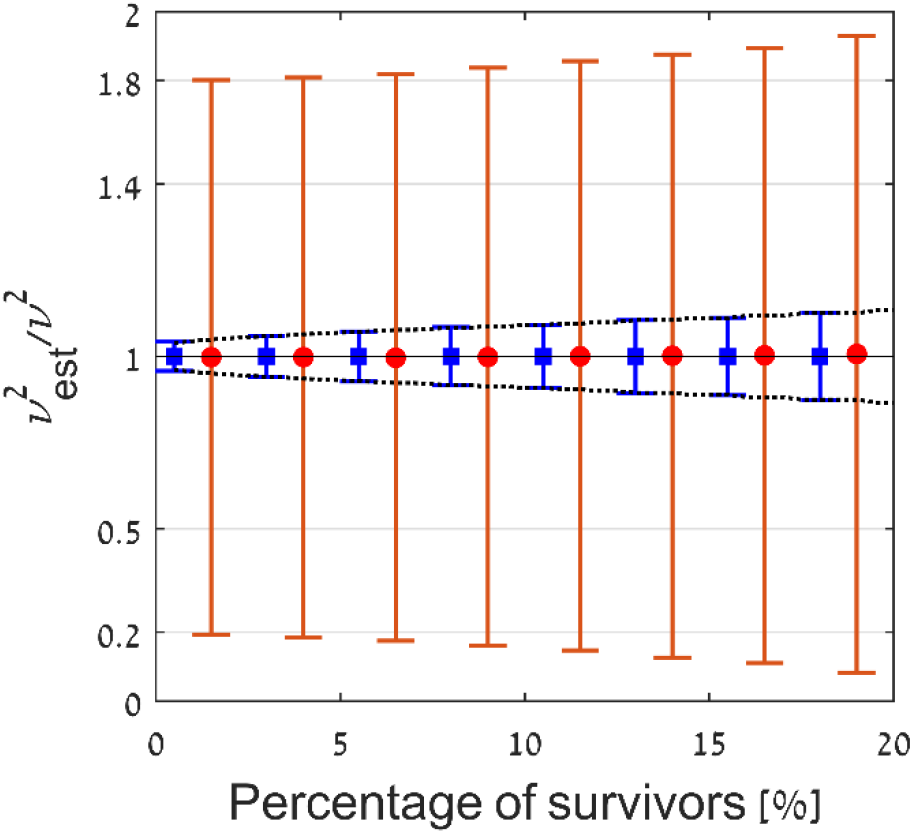
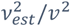 and 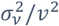 vs. the percentage of survivors. On the line *y* = 1, values of 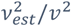, after averaging 10^6^ single-cell measurements by FFC (red circles), and PFC (blue squares). An exact result is obtained, without bias. For each value, the estimates of scaled variance of a single cell, ν, are also displayed. The variance is significantly smaller for PFC. The values of the variance obtained correspond exactly to the variance calculated in Eq. (11), for the method of comparison to an adjacent frame (dashed black line), and by Eq. (8), for the method of comparison to the first frame, for *p_limit_* approaching 0.

### Error Resulting from Unwanted Noise

#### Shot Noise

Of the expected disturbances in the experiment, Rothenberg found the most dominant noise was the Shot-noise: fluctuations due to variations in the number of photons emitted by the fluorophore. The noise variance is equal to the average signal and, for a single frame, is given by 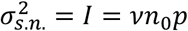. The photobleaching variance is 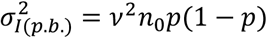. Therefore, the signal to noise ratio is 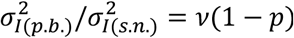. Because *ν* >> 1 (actually, the values *ν* and p can be controlled by frame exposure time or lighting intensity), the effect of shot-noise can be neglected. When calculating the photobleaching noise using Eq. (10), we use values from two measurements (*I_j-1_, I_j_* using PFC, or *I*_0_, *I_j_* using FFC). Each measurement contains a self-shot-noise, since subtracting values containing noises adds the noises together. For the FFC method the value of the shot-noise in frame *j* is 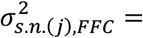 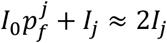 (The multiplication 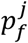 results from the same multiplication as in Eq. (10)). The final error in calculating *ν* is caused by shot-noises obtained after inserting 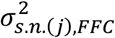 instead of 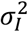 to Eq. (7). The resulting expression is:

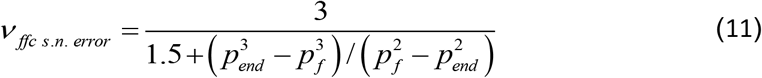

Similarly, in the PFC method, the final error is:

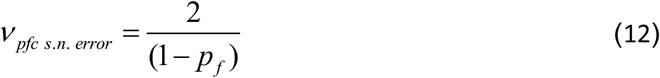

Because the S.N.R is given by 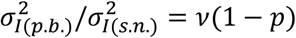, the noise is more dominant in the PFC method. this is because this method uses a constant *p_f_*, greater than the average 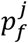, used in the FFC method. This result is understandable since the signal size is smaller when compared to a previous frame.

#### Background Noise

Each fluorescent image contains a background illumination, that does not result from the fluorophores. Rotenberg proposed to deal with this by reduction it, but the background illumination also has a shot-noise that remains after the background is removed. To analyze this, we express background illumination in the sampled area, for the purpose of measuring *I*_0_, as *I_b.g._*, and assume that *I_b.g._* remains constant throughout the measurement. The contribution to the variance in the fluorescent resulting from the background is 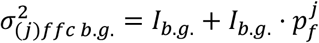 for FFC, and 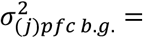 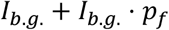 for PFC. To get the final error in *ν*, caused by background-noise we define the ratio between the background and the signal as *β*, i.e., *I_b.g._* = *βI*_0_. For FFC, 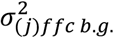 can be written into Eq. (7) (in place of 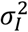). The resulting expression is:

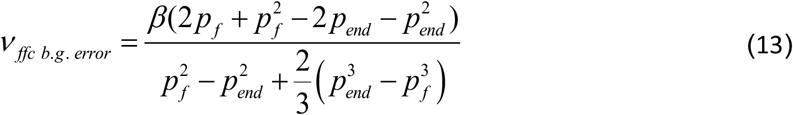

In PFC, the final error in *ν* is given by:

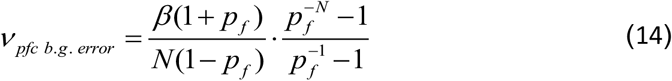

We also expect a greater background-noise effect in PFC.

The intensity of the signal depends on the number and type ([22]) of fluorophores measured. The background intensity depends on the type of substrate (for example, for LB substrate the background is stronger than for M9), and on its purity from fluorescence molecules. If the background illumination is due to fluorescence molecules undergoing photobleaching, then the background decays as a function of time, and causes unwanted photobleaching. The substrate molecules have their own decay constant and *ν*, and their noise cannot be corrected without knowing the *ν* value of the sample molecules.

#### Unwanted Noise Error Correction

In our experiment, the measuring signal itself is a variance therefore, in contrast to the normal effect of noise on experimental results, the noises discussed do not affect the result at random, but increase the value of the measured variance 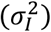. For this reason, these noises cannot be lowered by averaging many measurements, but the error can be corrected by estimating their size and subtracting them. In view of the above analysis, the measurement’s variance is 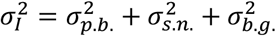, so we can correct the effect of the unwanted noise in Eq. (5) - by using 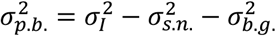 instead of 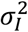:

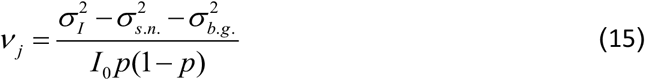

To test the effect of the noises, and the possibility of correcting them according to Eq. (17), we performed a simulation with shot-noise 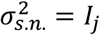 added for each *I_j_* value (Fig. 3.A), and its correction (Fig. 3.B); then, we performed a simulation with only background-noise 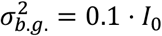 added for each *I_j_* value (Fig. 3.C) and its correction (Fig. 3.D). Finally, the two noises are added (3.E), and their corrections (3.F). The theoretical calculation of the noises for PFC (black dashed line) and for FFC (red dashed line) are also displayed. it can be seen that Eq. (13), (14), (15), and (16) accurately predict the errors due to shot-noise and background-noise, and allow accurate correction of the result. The calculation is done on 100 cells and for the purpose of averaging 10^5^ repetitions are made. the noise correction does not lead to an increase in standard deviation. Also, the effect of the noises is greater in PFC compared to FFC, and that the signal to noise ratio decreases as *ν* increases.

**Figure 3:**
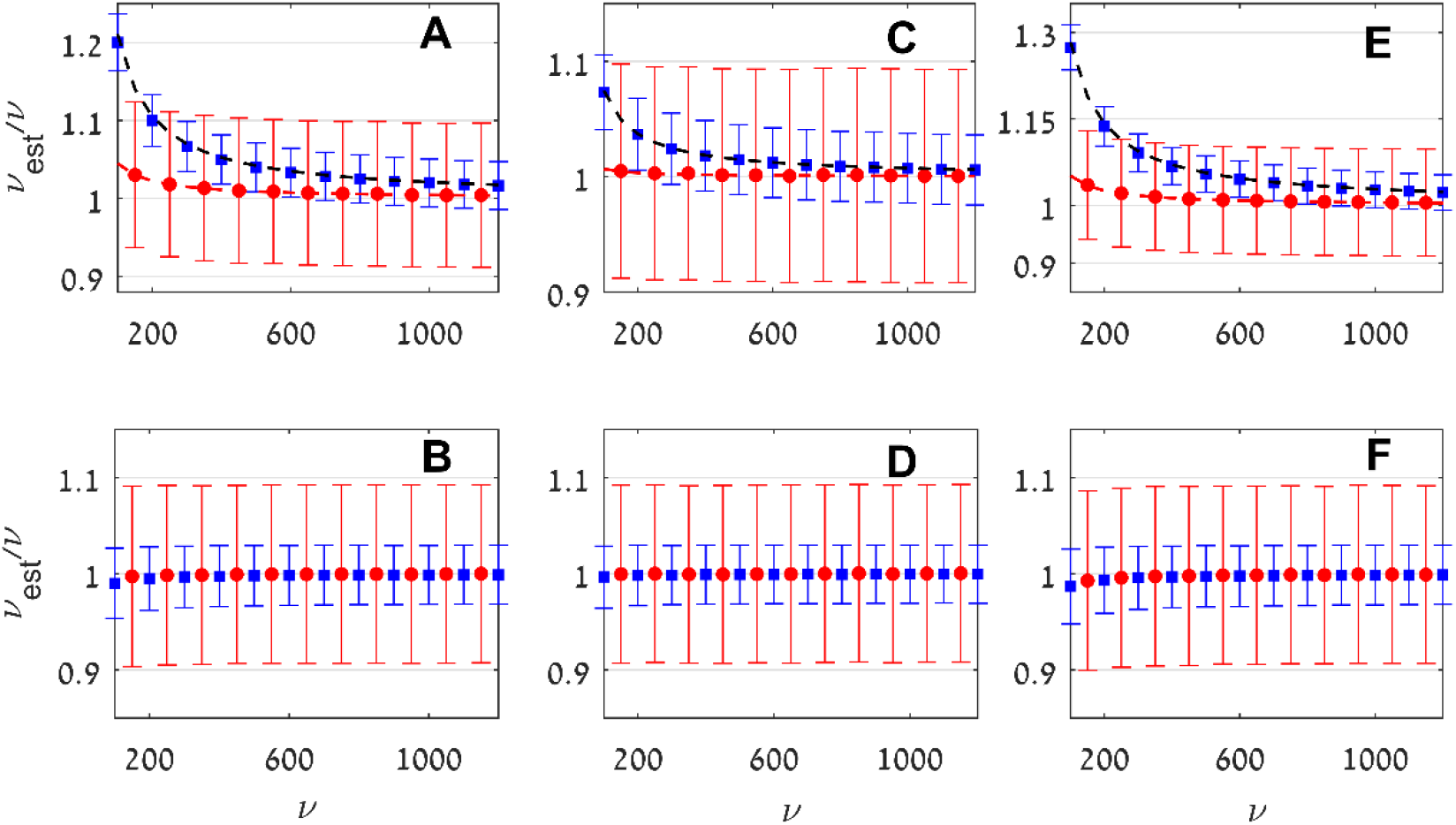
Simulation results showing ν*_est_*/ν vs ν. For all graphs, blue squares represent the result in PFC, with standard deviation; red circles – FFC with standard deviation. (A) Shot-noise 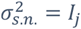 is added to each *I_j_* value. black dashed line: theoretical calculation by Eq. (14). Red dashed line: theoretical calculation by Eq. (13). (B) As in A, with noise correction. (C) Background-noise 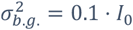 is added to each *I_j_* value. Black dashed line: theoretical calculation by Eq. (16). Red dashed line: theoretical calculation by Eq. (15). (D) As C with noise correction. (E) The two noises added. (F) As in E with correction for both noises. We used *n*_0_ = 1000, *p* = 0.9. The calculation is done on 100 cells and 10^5^ repeats for averaging. *p_f_* is calculated from simulation values in each experiment.

### Error from *p_f_* Inaccuracy

The calculation of the decay constant, *p_f_*, of the specific fluorophore in the experimental system is the initial stage of the algorithm. *p_f_* is used for calculating the expected fluorescence value in the following frames. In relation to these, the variance values are calculated. So, an inaccuracy in the calculation of *p_f_* results in an increase in the measured variance and an overestimation of *ν*.

The expression for error in *ν* due to error in *p_f_* in PFC is given by Eq. (18) (Sup. 2):

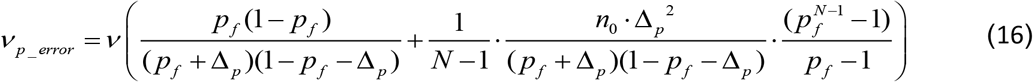

where *Δ_p_* is the error in *p_f_, ν_p_error_* is the error *ν*, estimated by using *p_f_* + *Δ_p_*, *n*_0_ number of initial fluorophores, and *N* is the number of frames.

For FFC, calculating *ν*_*p_error*_ is more complex because the expression is not integrable.

To compare the error from the two methods and testing Eq. (15), we performed a simulation where *p_f_*, with error *Δ_p_*, is used for the purpose of calculating *ν*.

It can be seen from Fig. (4) and Eq. (15) that the main contribution to the error is according to 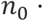 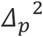, and Eq. (15) presents accurate predictions. Simulation results show that accuracy in determining *p_f_* is critical for obtaining reliable results, especially in cases where *n*_0_ is high. The FFC method is more susceptible to error in *p_f_* since it uses 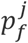, so the dominance of the error increases depending on *j*. The error in *p_f_* may be caused by the value of *p_f_* for each cell being slightly different. The main reason that may create this variance is uneven illumination in the sample plane. We discuss this in the next paragraph. in addition, a slight variability in the z axis position of the cells also leads to differences in illumination intensity between cells. However, whether the error in *p_f_* is positive or negative, the result is an increase in the measured variance, and therefore the error does not decrease following averaging over many cells.

**Figure 4:**
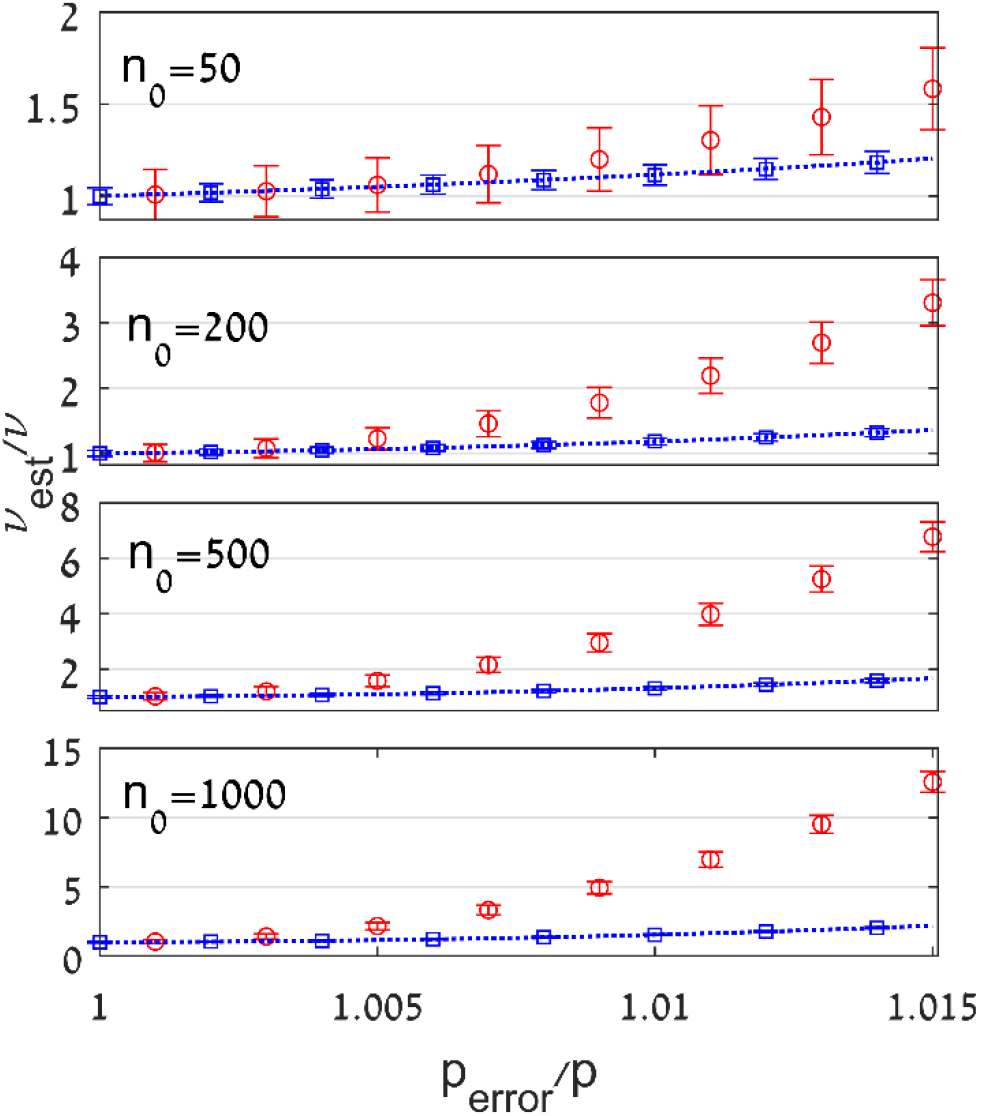
Relative error in ν vs. relative error in *p_f_* used for calculation, for *n*_0_ = 50, 200, 500, 1000, respectively. Blue squares: The result in PFC, including standard deviation. Red circles: FFC with standard deviation. Dashed blue Line: Result based on a theoretical calculation from Eq. (15). The results are obtained for averaging 10^4^ measurements, each with 50 cells, *p_end_* = 0.1, *p_f_* = 0.9.

By using an objective with a smaller Numerical Aperture (N.A.) it is possible to increase the depth of focus of the microscope, thus reducing the variance in illumination in the sample plane and in the z axis. Using a low N.A. leads to a larger spot size, and therefore, a lower illumination intensity and a smaller *ν*. This, on any case, can be compensated for by increasing exposure time. Another shortcoming in using a low N.A. is collecting the light at a smaller spatial angle, which also lead to a smaller *ν*. But, in this case, if we compensate with more collection time, then the photobleaching increases and leads to a smaller number of measurements for each cell.

If a calibration is made to v with a certain N.A., then the *ν* value for a different N.A. is obtained using *ν*_1_ = *ν*_2_(N. A._(1)_/N. A._(2)_)^4^. This is because two parameters that define *ν*, the illumination intensity and the spatial angle of the collection, depend on (N. A.)^2^.

A different error that can occur in *p_f_* calculation is an over-fitting to the experimental data, when the amount of data used for average *p_f_* is low. In this case, the inaccuracy leads to a reduction in the measured variance. By increasing the number of cells in the experiment, the effect of this error can be neglected.

### Uneven Illumination

Variation in illumination causes differences in the rate of photobleaching, i.e., *p_f_* differs depending on location. Even if Flat-Field Correction ([24]) is applied to the film, i.e. the intensity is calibrated according to the position, by photographing a fluorescent substrate with a uniform concentration of fluorophores and adjusting a two-dimensional function to the intensity obtained, to be used for calibration, still the dependence of *p_f_* on the position does not change, since *p_f_* is a ratio, and if the two values are calibrated in the same way then the ratio between them does not change. In contrast, Flat-Field Correction calibrates *ν*, because if the calibration value is *α*_(*x,y*)_, then 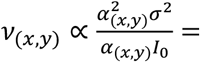 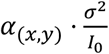, so that after performing the calibration the average *ν* can be used for cells at a different location.

If *p_f_* cannot be calculated from the population average, it is possible to calculate the decay constant for each cell by fitting an exponent function to its fluorescence data. In this way, however, there is a risk of over-fitting. If we use the values obtained from the fit as the expected values of the fluorescence, hence σ_i_ = *f*_(*i*)_ − *I_i_*, where *f_(i)_* is the fitted function, then an underestimation of the variance is created, because the fit minimizes the differences between the fluorescence values and the fitted function. This minimization leads to an error in v calculation, as shown in simulation results (Fig. 5, black stars). The error becomes smaller as a function of the number of frames. This results from the decreased effect of fitting process to reduce the variance. It is possible to use *f_(i)_* only for the purpose of calculating *p_f_* for every cell. Here, also, *p_f_* error is expected due to over-fitting. After calculating *p_f_*, FFC or PFC can be used for the quantification. Simulation results (blue squares) show that for a measurement with enough frames, PFC allows negligible error. In any case, this method makes it possible to set an upper limit for estimating the number of proteins. Performing the fitting requires knowledge of the decay, and an appropriate function must be used if the fluorescence decays like the bi-exponent model (Sup. 3), as detailed in the next paragraph.

**Figure 5:**
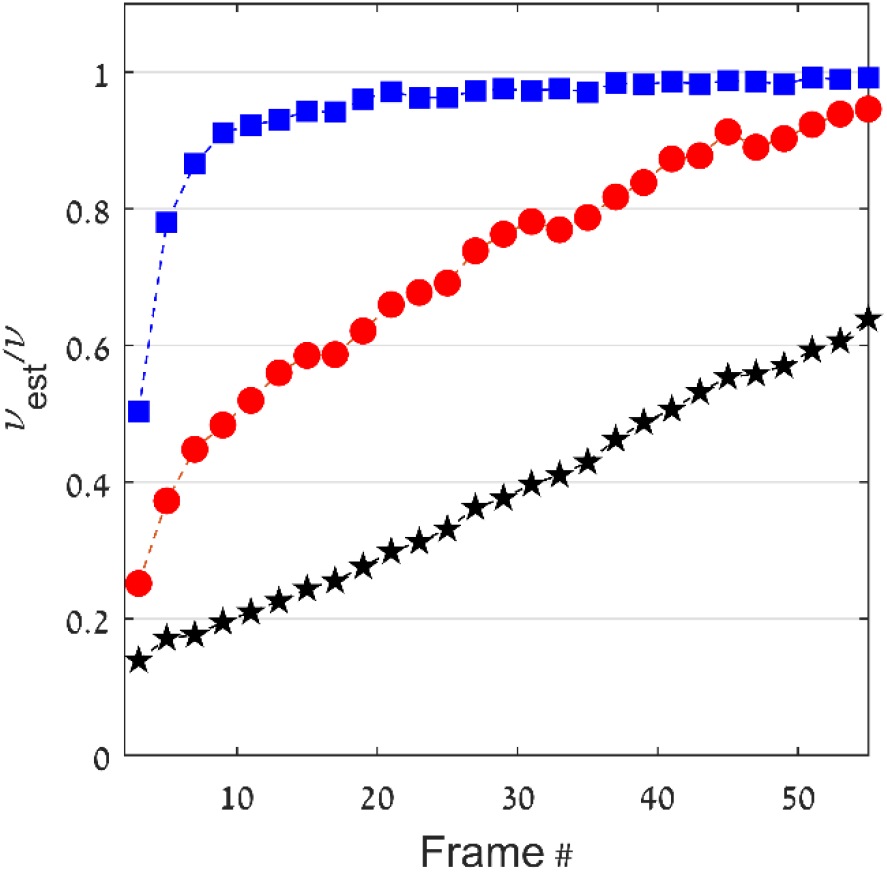
Calculating ν*_est_*/ν by fitting a function of a decaying exponent to each cell individually, where the fitting is used for *p_f_* calculation only, and the quantification is done using PFC (blue squares) and FFC (red circles); and where the fitting values are used as the expected value of the fluorescence (black stars). Results are obtained for averaging 10^3^ cell measurements, and *p_f_* = 0.9.

The error resulting from over-fitting does not depend on *n*_0_. We have shown that the error resulting from *Δ_p_* is proportional to *n*_0_ · *Δ_p_*^2^, and that *Δ_p_* caused by over-fitting is proportional to the ratio between the standard deviation of the protein number to the mean protein number 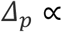 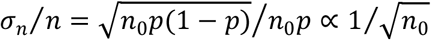, so *n*_0_ is reduced.

### Matching the Algorithm to the Bi-Exponential Decay Model

Recently, Peter S. Swain [19] showed that the decay of the fluorescence is non-exponential. So, a test was made for the adaptation of the algorithm to the bi-exponential decay model. Initially, a simulation is performed in which the *p_f_* value used to prepare the simulation data varies in each frame continuously, in the range between 0.85 and 0.95. After preparing the fluorescence data, 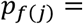 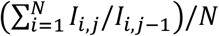 is calculated for each frame and the quantification is performed using Eq. (11) (PFC). In addition, 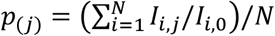 is calculated and quantification is performed with Eq. (10) (FFC). In this case, there is no error in calculating *ν* (data not shown). Another examined case is one with difference in decay between two groups of proteins in the same cell, such as cytoplasmic proteins and envelope proteins. For this purpose, a simulation is performed with each cell having half of the initial fluorophores, simulated with decay constant *p*_*f*,1_, and the other half with *p*_*f*,2_. *v* value are equal for both types of fluorophores. Figure (6) shows the results where *p*_*f*,1_ = 0.85 and *p*_*f*,2_ are determined in each experiment, in the range between 0.67 and 0.97.

**Figure 6:**
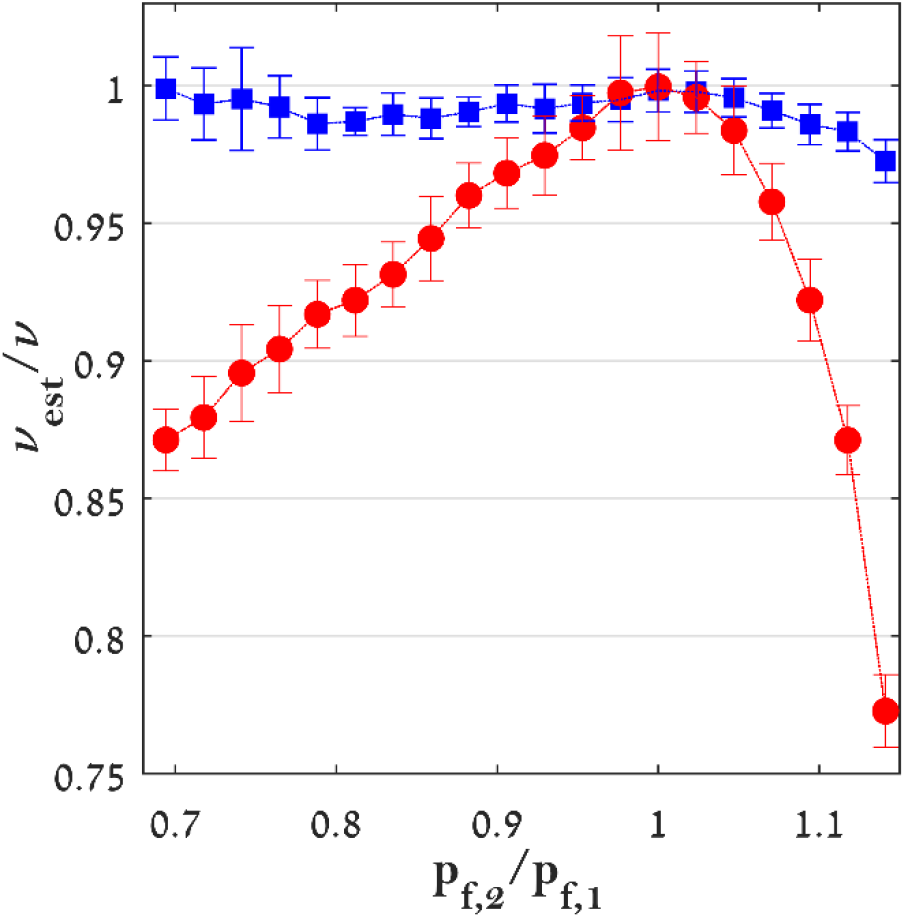
Relative error in ν vs *p*_*f*,2_/*p*_*f*,1_. Blue squares: estimate using PFC. Red circles: estimate using FFC. *p*_*f*,1_ is constant and equal to 0.85. The results were obtained for averaging 20 measurements each of which included 3000 cells, ν = 753. *n*_0_ = 1000.

When *p*_*f*,2_/*p*_*f*,1_ = 1, we get, as expected, an accurate calculation of *ν*. On the other hand, when there is a difference between the photobleaching rates, a negligible error is obtained for PFC, and a more significant one for FFC. The magnitude of the error generated is asymmetrical with respect to *p*_*f*,2_/*p*_*f*,1_ = 1, because the error resulting from the use of an incorrect *p* is determined by 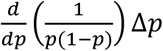, and has a minimum for *p* = 0.5 and an extremum for *p* = 1 and *p* = 0. So, when *p*_*f*,2_ tends to 1 the error increases (Sup. 4).

The third case examined is when there is a slight cell-to-cell variance in *p_f_* values. In this situation, the average *p_f_* calculation does not express the specific decay constant of each cell. Therefore, there may be a significant *ν* calculation error, as described in the *p* error effect analysis. It follows that if the changes in decay rates throughout the experiment are the same for all cells, the algorithm is still able to provide reliable results. When there is a difference between the cells themselves, on the other hand, the inability to determine the decay constant of a single cell can lead to a significant error.

### Estimate Using Gaussian Process

Recently Peter S. Swain [19] used a technique similar to the one developed by Rotenberg for protein quantification (his basic Eq. [6 in his article] is the same as Eq. 5), except that they calculate the expected value of fluorescence using the Gaussian Process. He verifies his approach using six different proteins tagged with Green Fluorescent Protein (GFP) in budding yeast and compare their results to estimates made using biochemical techniques, such as quantitative Western blotting and mass spectrometry. He showed that there is a large variance between the decay constants of the different cells. The difference in the decay constants may be explained by the uneven illumination of the cells – the data for the experiments is taken from regions where the illumination intensity is at least 80% of the median illumination. That is, there is a large variability in illumination to which the cells are exposed, and as explained above, flat field correction does not calibrate the decay constants (*p* values) created due to uneven illumination.

The curve obtained by the Gaussian Process is not limited to being “smooth”. Its flexibility is determined by three hyperparameters for a squared exponential Gaussian Process. In our case, these parameters are of special importance because they may cause over-fitting, reducing measured variance. Fit flexibility is expressed through the change in *p_f_* values throughout the experiment. Fig. (7) shows *p_f_* values calculated as the ratio of the fit values in adjacent frames, obtained using GP (according to the original data, and algorithm, attached to Peter’s article). This flexibility does not represent a physical phenomenon, and leads to reduced measured variability and overestimation.

**Figure 7:**
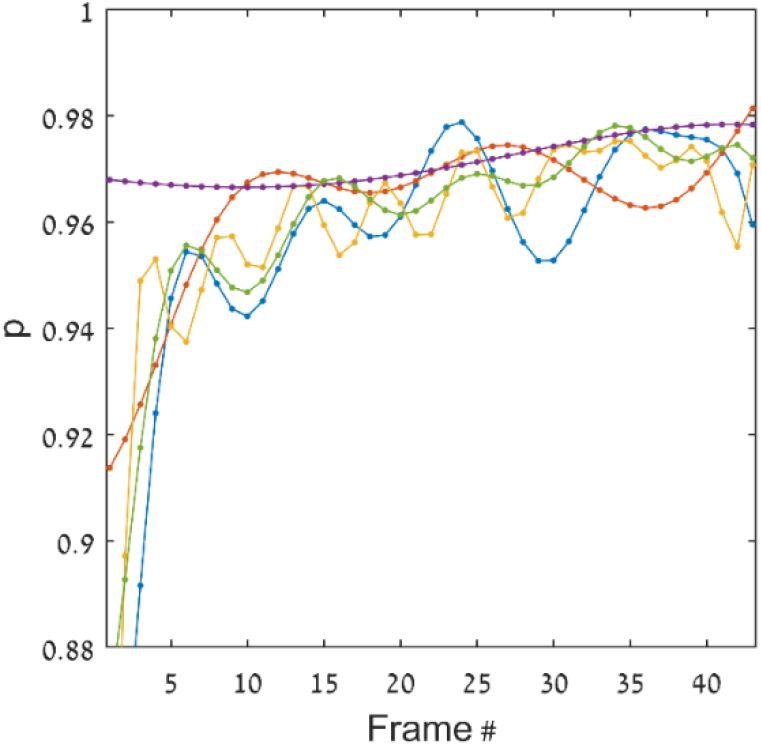
The ratio of the signal of two adjacent frames estimated using GP. The figure shows data from the first five cells in the first experiment (Fus3) from Peter’s data.

To test the effect of different experimental conditions on the quantification, we used the measurement data, and the original code, to process GP, on Peter’s results. In the first stage, an accurate reconstruction of the results presented in Peter’s article was performed. In the second stage, we added a sum of the two illumination values from adjacent frames, to simulate experimental data with half the frames, each frame with double exposure time. Because *ν* and *p_f_* are proportional to exposure time, we get experimental data with *ν* values approximately doubled, and *p_f_* values increased by a power of two. Similarly, all four measurement values are added for the purpose of imaging an experimental system in which *ν* value is approximately quadruple and *p_f_* increased by a power of four. The median values obtained for quantification of each protein are shown in Fig. (8), in comparison with the quantification results obtained from biochemical methods, as presented in Peter’s article. It can be seen that changing the conditions has a significant impact on the results.

**Figure 8:**
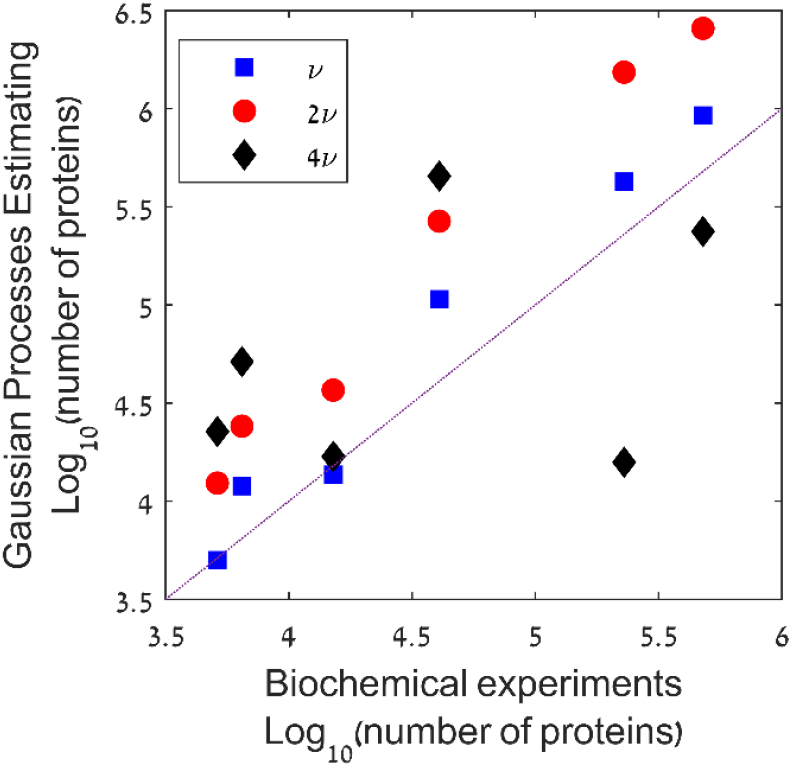
estimates made using Gaussian Process vs. estimates made using biochemical techniques. Blue square: estimates from swain’s original data. Red circles: estimates for 2ν. Black rhombus: estimates for 4ν.

It is evident that increasing *ν* increases the estimate of proteins’ number (except in three cases for 4*ν*). This result is expected because from the analysis of the results it is obtained that *ν* ≈ 20 And as explained, the ratio between the photobleaching-noise and the shot-noise is given by *ν* (1-p). Since *p* ≈ 0.96, it is observed that the shot-noise has a large effect (no data is given to allow the background-noise to be evaluated). This leads to underestimation, and the effect gets smaller as *ν* increases.

## DISCUSSION AND CONCLUSIONS

Quantifying the number of events by calculating the ratio of the fluctuations to the signal average is a useful and proven research tool for random processes with a fixed average. Rotenberg first proposed applying this technique for processes with the mean signal decaying as a function of time.

Because of the variance in the initial protein number of the cells, the value around which the fluorescence of a single cell is expected to be distributed is not the average fluorescence of the cells but should be calculated according to the initial fluorescence of that cell and the average decay of the whole population.

The variance of the measurement can be reduced by calculating the expected value of the fluorescence in a particular frame, relative to the previous frame, not to the first frame of the cell. In this case, there is no correlation between the random events from the different frames used to calculate *ν*, and therefore the variance is minimal. The disadvantage of this method is that the dominance of the unwanted noises is greater. On the other hand, the method of comparison to the previous frame is less sensitive to the effect of error in calculating *p*, which may result from different cell illumination conditions, and less sensitive to error due to variability in fluorophore decay rates.

The dominant unwanted noises in the experiment are shot-noise and background-noise. But because the measurement in the experiment is noise measurement, the unwanted noises are not randomly included in the final result, but as an expected and evaluable value. Because of this, we can subtract these noises and get the effect of the photobleaching noise only. In any case, as Rotenberg showed, designing an experiment with high *ν* values leads to noise reduction, so its effect can be neglected.

In performing the experiment there are several challenges, which without attention can lead to a significant error. Uneven illumination or uneven light collection leads to dependence on the decaying constants and *ν* values on location. Calibrating the image using flat field correlation calibrates the *ν* values, but not the decay constants. It seems that the most significant condition for the success of an experiment is the possibility of calculating the decay constant with the required accuracy, since even a small error in the decay constant calculated can cause a significant error in the calculated ν. But the variance in the decay constant does not allow calculating its mean value from deferent cells. If equal illumination conditions cannot be assumed for all cells, it is possible to calculate a relatively close lower limit for the number of proteins, by calculating the decay of each cell from its decay curve only.

The algorithm can be adjusted for cases where the decay rate of the fluorescence is not constant, if the rate changes are common to all cells. In case there are several groups of fluorophores with different decay rates, the algorithm leads to negligible error for PFC, and a larger error for FFC.

Analysis of photobleaching fluctuations provides a straightforward quantification of proteins number, and as Rotenberg wrote, can be applied to any superposition of *n*_0_ discrete decaying processes. However, the evaluation of the expected errors in the quantification is essential for planning the experimental conditions, and for evaluating the error.

## BIBLIOGRAPHY

[1] D. R. Spiegel and R. J. Helmer, “Shot‐noise measurements of the electron charge: An undergraduate experiment”, Am. J. Phys., 1995, doi: 10.1119/1.17867.

[2] R. de-Picciotto, M. Reznikov, M. Heiblum, V. Umansky, G. Bunin, and D. Mahalu, “Direct observation of a fractional charge”, Nature, vol. 389, no. 6647, pp. 162–164, Sep. 1997, doi: 10.1038/38241.

[3] L. Saminadayar, D. C. Glattli, Y. Jin, and B. Etienne, “Observation of the e/3 Fractionally Charged Laughlin Quasiparticle”, Phys. Rev. Lett., vol. 79, no. 13, pp. 2526–2529, Sep. 1997, doi: 10.1103/PhysRevLett.79.2526.

[4] N. Rosenfeld, T. J. Perkins, U. Alon, M. B. Elowitz, and P. S. Swain, “A Fluctuation Method to Quantify In Vivo Fluorescence Data”, Biophys. J., vol. 91, no. 2, pp. 759–766, 2006, doi: 10.1529/biophysj.105.073098.

[5] S.W. Teng et al., “Measurement of the Copy Number of the Master Quorum-Sensing Regulator of a Bacterial Cell”, Biophys. J., vol. 98, no. 9, pp. 2024–2031, May 2010, doi: 10.1016/j.bpj.2010.01.031.

[6] L. Zamparo and T. J. Perkins, “Statistical lower bounds on protein copy number from fluorescence expression images”, Bioinformatics, vol. 25, no. 20, pp. 2670–2676, Oct. 2009, doi: 10.1093/bioinformatics/btp415.

[7] V. C. Coffman and J.Q. Wu, “Counting protein molecules using quantitative fluorescence microscopy”, Trends Biochem. Sci., vol. 37, no. 11, pp. 499–506, 2012, doi: 10.1016/j.tibs.2012.08.002.

[8] H. G. Garcia, H. J. Lee, J. Q. Boedicker, and R. Phillips, “Comparison and Calibration of Different Reporters for Quantitative Analysis of Gene Expression”, Biophys. J., vol. 101, no. 3, pp. 535– 544, 2011, doi: 10.1016/j.bpj.2011.06.026.

[9] X. Wang, S. Wang, and H. Ma, “Characterization of local polarity and structure of Cys121 domain in β-lactoglobulin with a new thiol-specific fluorescent probe”, Analyst, vol. 133, no. 4, p. 478, Apr. 2008, doi: 10.1039/b717230c.

[10] R. Wang, C. Yu, F. Yu, L. Chen, and C. Yu, “Molecular fluorescent probes for monitoring pH changes in living cells”, TrAC Trends Anal. Chem., vol. 29, no. 9, pp. 1004–1013, Oct. 2010, doi:10.1016/J.TRAC.2010.05.005.

[11] K. L. Budzinski, M. Zeigler, B. S. Fujimoto, S. M. Bajjalieh, and D. T. Chiu, “Measurements of the Acidification Kinetics of Single SynaptopHluorin Vesicles”, Biophys. J., vol. 101, no. 7, pp. 1580– 1589, Oct. 2011, doi: 10.1016/j.bpj.2011.08.032.

[12] F. Ye, C. Wu, Y. Jin, Y.-H. Chan, X. Zhang, and D. T. Chiu, “Ratiometric Temperature Sensing with Semiconducting Polymer Dots”, J. Am. Chem. Soc., vol. 133, no. 21, pp. 8146–8149, Jun. 2011, doi: 10.1021/ja202945g.

[13] F. Li, A. H. Westphal, A. T. M. Marcelis, E. J. R. Sudhölter, M. A. Cohen Stuart, and F. A. M. Leermakers, “Thermally sensitive dual fluorescent polymeric micelles for probing cell properties”, Soft Matter, vol. 7, no. 23, p. 11211, Nov. 2011, doi: 10.1039/c1sm06597a.

[14] C. R. Nayak and A. D. Rutenberg, “Quantification of Fluorophore Copy Number from Intrinsic Fluctuations during Fluorescence Photobleaching”, Biophys. J., vol. 101, no. 9, pp. 2284–2293, 2011, doi: 10.1016/j.bpj.2011.09.032.

[15] A. T. Lombardo et al., “Myosin Va molecular motors manoeuvre liposome cargo through suspended actin filament intersections in vitro”, Nat. Commun., vol. 8, p. 15692, 2017, doi: 10.1038/ncomms15692.

[16] S. R. Nelson, K. M. Trybus, and D. M. Warshaw, “Motor coupling through lipid membranes enhances transport velocities for ensembles of myosin Va”, Proc. Natl. Acad. Sci., vol. 111, no. 38, pp. E3986–E3995, 2014, doi: 10.1073/pnas.1406535111.

[17] N. H. Kim et al., “Real-time transposable element activity in individual live cells”, doi: 10.1073/pnas.1601833113.

[18] E. Crozat et al., “Post-replicative pairing of sister ter regions in Escherichia coli involves multiple activities of MatP”, Nat. Commun., vol. 11, no. 1, pp. 1–12, 2020, doi: 10.1038/s41467-020-17606-6.

[19] E. Bakker and P. S. Swain, “Estimating numbers of intracellular molecules through analysing fluctuations in photobleaching”, Sci. Rep., vol. 9, no. 1, Dec. 2019, doi: 10.1038/s41598-019-50921-7.

[20] P. S. Swain et al., “Inferring time derivatives including cell growth rates using Gaussian processes”, Nat. Commun., vol. 7, no. 1, pp. 1–8, Dec. 2016, doi: 10.1038/ncomms13766.

[21] Y. Taniguchi et al., “Quantifying E. coli Proteome and Transcriptome with Single-Molecule Sensitivity in Single Cells”, Science (80-.)., vol. 329, no. 5991, pp. 533–538, Jul. 2010, doi: 10.1126/science.1188308.

[22] G. Cannone, F., Caccia, M., Bologna, S., Diaspro, A. and Chirico, “Single molecule spectroscopic characterization of GFP‐mut2 mutant for two‐photon microscopy applications”, Microsc. Res. Tech., vol. 65, no. 4–5, 2004.

[23] N. H. Kim, G. Lee, N. A. Sherer, K. M. Martini, N. Goldenfeld, and T. E. Kuhlman, “Real-time transposable element activity in individual live cells”, Proc. Natl. Acad. Sci., vol. 113, no. 26, pp. 7278–7283, 2016, doi: 10.1073/pnas.1601833113.

[24] J. C. Waters and T. Wittmann, “Concepts in quantitative fluorescence microscopy”, Quant. Imaging Cell Biol., vol. 123, pp. 1–18, 2014, doi: 10.1016/B978-0-12-420138-5.00001-X.

