## Supplementary material for "Variance Reducing and Noise Correction in Protein Quantification by Measuring Fluctuations in Fluorescence due to Photobleaching": sup.

SUPPLEMENTARY MATERIALS

### 1. Development of the expression for an error caused by a difference in the values of the initial fluorophore's according to the Rotenberg method

$\alpha$ is defined such that $\langle I_0\rangle = I_{(0,i)}(1+\alpha_i)$. Since $\langle I_j\rangle = \langle I_0\rangle p$, the expression for variance is:

$$\sigma^2_{I(j,i)} = \left(I_j - \langle I_j\rangle\right)^2 = \left(\langle I_0\rangle p + \alpha\langle I_0\rangle p - \langle I_j\rangle\right)^2 = \left(\sigma_{I(j,i)} + \alpha\langle I_0\rangle p\right)^2$$
$$= \sigma^2_{I(j,i)} + (\alpha\langle I_0\rangle p)^2 + 2\sigma_{I(j,i)}\cdot\alpha\langle I_0\rangle p$$
$$\sigma^2_{I(j,i)} = \left(I_{j,i} - \langle I_j\rangle\right)^2 = \left(I_{j,i} - \langle I_0\rangle p\right)^2 = \left(I_{j,i} - I_{0,i}(1+\alpha_i)p\right)^2 = \left(\sigma_{I(j,i)} + \alpha I_{0,i}p\right)^2$$
$$= \sigma^2_{I(j,i)} + \left(\alpha I_{0,i}p\right)^2 + 2\sigma_{I(j,i)}\cdot\alpha I_{0,i}p$$

Since $\left\langle\sigma_{I(j,i)}\cdot\alpha I_0 p\right\rangle = 0$, the error is $\sigma^2_{error} = (\alpha\langle I_0\rangle p)^2$, so the signal to noise ratio is:

$$\frac{I_0\nu p(1-p)}{(\alpha\langle I_0\rangle p)^2} = \frac{1-p}{\alpha^2 n_0 p}$$

The contribution of the error in the calculation of the variance to the final calculation of $\nu$, according to Eq. (7), can be calculated like so:

$$\nu_{error} = \frac{\int_{p_f}^{p_{end}}\sigma^2_{error}dp}{I_0\int_{p_f}^{p_{end}}p(1-p)dp} = \alpha^2\langle I_0\rangle/\left((1.5p^2_{end} - 1.5p_f^2)/(p^3_{end} - p_f^3) - 1\right)$$

The reason the integral starts from $p_f$ and not from 0 is that, in practice, the integral is not continuous, but is calculated using the trapezoidal method. In this method each frame has a contribution to the integral amount except for the first one ($j = 0$), since no noise can be calculated for it in relation to what is expected from a previous frame.

### 2. Development of an equation for error in $\nu$ due to error in $p$

$$\nu = \frac{\sigma^2}{I_n p(1-p)} = \frac{\left(I_j p - I_{j-1}\right)^2}{I_n p(1-p)} =$$

$$\nu_{p+\Delta_p} = \frac{\left(I_j(p+\Delta_p) - I_{j-1}\right)^2}{I_j(p+\Delta_p)(1-p-\Delta_p)} = \frac{\left(I_j p + I_j\Delta_p - I_{j-1}\right)^2}{I_j(p+\Delta_p)(1-p-\Delta_p)} = \frac{\left(\sigma + I_j\Delta_p\right)^2}{I_j(p+\Delta_p)(1-p-\Delta_p)}$$

$$= \frac{\sigma^2 + 2\sigma I_j\Delta_p + I_j^{\,2}{\Delta_p}^2}{I_j(p+\Delta_p)(1-p-\Delta_p)} = \nu\frac{p(1-p)}{(p+\Delta_p)(1-p-\Delta_p)} + \frac{2\sigma\Delta_p + I_j{\Delta_p}^2}{(p+\Delta_p)(1-p-\Delta_p)}$$

$$E(2\sigma\Delta_p) = 0$$

$$\Rightarrow \nu_{p+\Delta_p} = \nu\frac{p(1-p)}{(p+\Delta_p)(1-p-\Delta_p)} + \frac{I_j{\Delta_p}^2}{(p+\Delta_p)(1-p-\Delta_p)}$$

$$= \nu\left(\frac{p(1-p)}{(p+\Delta_p)(1-p-\Delta_p)} + \frac{n_j{\Delta_p}^2}{(p+\Delta_p)(1-p-\Delta_p)}\right)$$

This is the expression for $\nu$ with the error as calculated from a pair of adjacent frames, in practice the N is calculated from the average of the results from all the frames, where N is the number of frames then the number of measurements is N-1

$$\nu_{mean,p+\triangle p} = \frac{1}{N-1}\sum_{j=0}^{j=N-2}\nu\left(\frac{p(1-p)}{(p+\Delta_p)(1-p-\Delta_p)} + \frac{n_0(p)^j{\Delta_p}^2}{(p+\Delta_p)(1-p-\Delta_p)}\right)$$

$$= \nu\frac{p(1-p)}{(p+\Delta_p)(1-p-\Delta_p)} + \frac{\nu}{N-1}\sum_{j=0}^{j=N-2}\left(\frac{n_0(p)^j{\Delta_p}^2}{(p+\Delta_p)(1-p-\Delta_p)}\right)$$

$$= \nu\frac{p(1-p)}{(p+\Delta_p)(1-p-\Delta_p)} + \frac{\nu}{N-1}\cdot\frac{n_0\cdot{\Delta_p}^2}{(p+\Delta_p)(1-p-\Delta_p)}\sum_{j=0}^{j=N-2}(p)^j$$

$$= \nu\left(\frac{p(1-p)}{(p+\Delta_p)(1-p-\Delta_p)} + \frac{1}{N-1}\cdot\frac{n_0\cdot{\Delta_p}^2}{(p+\Delta_p)(1-p-\Delta_p)}\cdot\frac{(p^{N-1}-1)}{p-1}\right)$$

### 3. Calculating $p_f$ by fitting a function of a two-decaying exponent to each cell individually

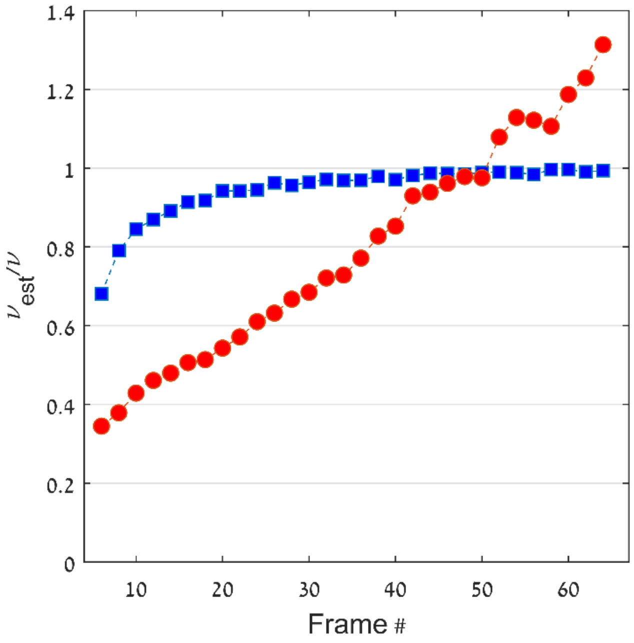


*Calculating $\nu_{est}/\nu$ by fitting a function of a two-decaying exponent to each cell individually, where the fitting is used for $p_f$ calculation only, and the quantification is done using PFC (blue squares) and FFC (red circles); Results are obtained for averaging $10^3$ cell measurements, and $p_f$ var between 0.8-0.9.*

### 4. Estimate for cells with two decaying rate fluorophores

It can be seen from the figure that the error in $P$ increases as it approaches 1.

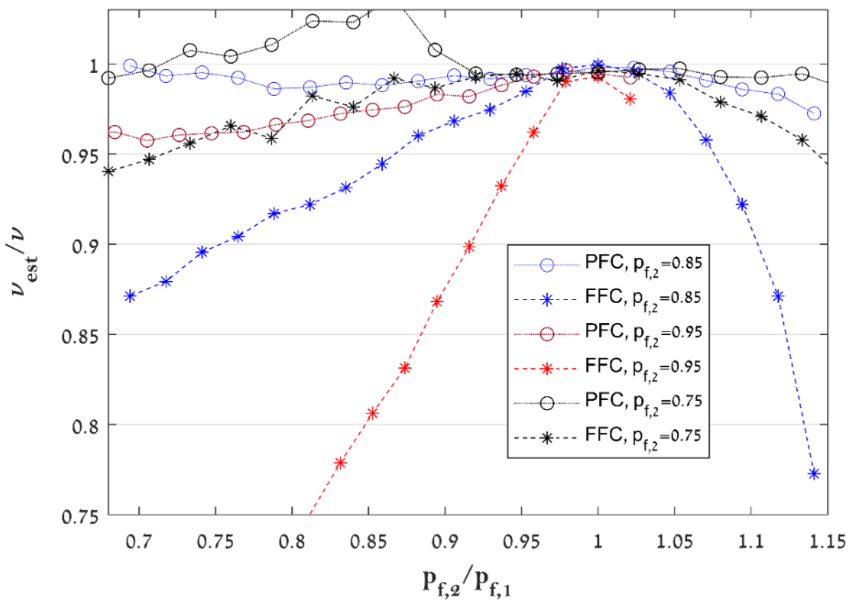


*Figure 1: Relative error in $\nu$ vs $p_{f,2}/p_{f,1}$. Circles: estimate using PFC. Stars: estimate using FFC. blue lines: $p_{f,1} = 0.85$ constant. Red lines: $p_{f,1} = 0.75$ constant. Black lines: $p_{f,1} = 0.95$ constant. The results were obtained by averaging over 20 measurements, each of which included 3000 cells, $\nu = 753. n_0 = 1000$.*
